# Systemic Markers of Adaptive and Innate Immunity Are Associated with COPD Severity and Spirometric Disease Progression

**DOI:** 10.1101/193920

**Authors:** Eitan Halper-Stromberg, Jeong H. Yun, Margaret M. Parker, Ruth Tal Singer, Amit Gaggar, Edwin K. Silverman, Sonia Leach, Russell P. Bowler, Peter J. Castaldi

**Author notes:** **Corresponding Authors**: Peter Castaldi, MD, MSc, Channing Division of Network Medicine, Brigham and Women's Hospital, Boston, MA 02115, Russell P. Bowler, MD, PhD, National Jewish Health, Denver, CO. authors contributed equally (Russell P. Bowler, Peter J. Castaldi). **Abbreviations** AUC: area under the curve CBC: complete blood count COPD: chronic obstructive pulmonary disease CT: computed tomography FEV_1_: forced expiratory volume in one second GOLD: Global Initiative for Obstructive Lung Disease LAA950: low attenuation area less than 950 Hounsfield units (computed tomorgraphy based emphysema measure) LR: linear regression NBS: network-based stratification ROC: receiver operating characteristic SVM: support vector machine.

## Abstract

The progression of chronic obstructive pulmonary disease (COPD) is associated with marked alterations in circulating immune cell populations, but no studies have characterized alterations in these cell types across the full spectrum of lung function impairment in current and former smokers. In 6,299 subjects from the COPDGene and ECLIPSE studies, we related Coulter blood counts and proportions to cross-sectional FEV_1_ adjusting for current smoking status. We also related cell count measures to three-year change in FEV_1_ in ECLIPSE subjects. In a subset of subjects with blood gene expression data, we used cell type deconvolution methods to infer the proportions of immune cell subpopulations, and we related these to COPD clinical status. We observed that FEV_1_ levels are positively correlated with lymphocytes and negatively correlated with myeloid populations such as neutrophils and monocytes. In multivariate models, absolute cell counts and proportions were associated with cross-sectional FEV_1_, and lymphocytes, monocytes, and eosinophil counts were predictive of three-year change in lung function. Using cell type deconvolution to study immune cell subpopulations, we observed that subjects with COPD had a lower proportion of CD4+ resting memory cells and naive B cells compared to non-COPD smokers. Alterations in circulating immune cells in COPD support a mixed pattern of lymphocyte suppression and an enhanced myeloid cell immune response. Cell counts and proportions contribute independent information to models predicting lung function suggesting a critical role for immune response in long-term COPD outcomes. Cell type deconvolution is a promising method for immunophenotyping in large cohorts.

## Background

Chronic obstructive pulmonary disease (COPD) is associated with profound alterations in immune cells within the lung and in the systemic circulation. The systemic inflammation may reflect “spill-over” of inflammatory processes within the lung, primary alterations in the extra-pulmonary immune response, or a combination of both processes(1). These alterations affect cell types involved in both the innate(2-6) and adaptive(7-15) immune response. In population studies(16, 17) and a systematic review(18), total counts of peripheral leucocytes were associated with cross-sectional and prospective changes in lung function, but few studies have been performed in large cohorts with detailed cell count data to observe relationships across the full spectrum of lung function. In addition, current smoking has an independent effect on immune cells(8, 19) and often serves as an important potential confounder of immunologic studies of current and former smokers with COPD.

Cell type quantification by flow cytometry is rarely available from large, population-based studies of COPD. However, novel, cell-type “deconvolution” approaches have been shown to infer accurately the relative proportions of immune cell types from genome-wide blood gene expression data(20, 21). Thus, cell-type deconvolution is a potentially powerful approach to enable the simultaneous study of many different cell types in large cohorts of subjects with available blood gene expression, but it has not yet been applied to cohorts of subjects with COPD.

We hypothesized that 1) peripheral immune cell types quantified through Coulter complete blood counts (CBC) have significant associations to cross-sectional FEV_1_ and prospective FEV_1_ decline; and that 2) cell-type deconvolution methods can enable the simultaneous study of multiple immune cell subpopulations in cohorts of smokers with COPD and blood gene expression data. We explored the first hypothesis in two large cohorts of smokers enriched for COPD, the COPDGene and ECLIPSE studies, which enabled the characterization of immune cell profiles across the full spectrum of lung function impairment while accounting for current smoking effects. The large number of study subjects allowed for detailed modeling of the relationship between multiple cell types, current smoking status, and lung function. To explore the second hypothesis, we used two cell type deconvolution methods to infer immune cell subpopulation proportions in a subset of smokers with blood gene expression data in the ECLIPSE Study, and we validated these inferred cell type proportions against measured CBC data. We then compared levels of inferred circulating immune cell subpopulations by COPD status, confirming that inferred estimates of circulating immune cell types such as monocytes, naive B cells, and resting T memory cells are altered in the COPD state.

## Methods

### Study Populations

Recruitment criteria and study protocols for the ECLIPSE and COPDGene studies have been previously reported. COPDGene enrolled 10,192 subjects across the entire GOLD spectrum between the ages of 45 and 80 with at least a 10 pack-year smoking history(22). These subjects completed their Phase 1 study visit between 2007-2011. As of September 24, 2016, 5000 subjects had completed their Phase 2 five-year follow-up visit which included all of the data items collected in Phase 1 as well as complete blood count data, which was not collected at the Phase 1 visit.

The Evaluation of COPD Longitudinally to Identify Predictive Surrogate Endpoints (ECLIPSE) study was a multicenter study that enrolled subjects aged 40-75 with COPD and at least a 10-year smoking history (COPD defined by FEV_1_< 80% of predicted and FEV_1_/FVC <= 0.7) or who were smokers without COPD (FEV_1_>85% and FEV_1_/FVC > 0.7). Details of this study have been previously published(23). Gene expression analyses were performed in a subset of subjects in this study from whom genome-wide gene expression data were generated on the Affymetrix Human U133 Plus2 chip as previously reported(24). For both COPDGene and ECLIPSE, the institutional review boards of all participating centers approved these studies, and written informed consent was obtained from all subjects.

### Phenotype and Covariate Definitions

In COPDGene and ECLIPSE, spirometry was performed before and after administration of 180 mcg of albuterol according to international guidelines(25). COPD cases and GOLD stages were defined according to GOLD spirometric criteria (FEV1 % of predicted < 80% and FEV1/FVC < 0.7) (26). Subjects with preserved ratio impaired spirometry (PRISm) were defined by post-bronchodilator FEV_1_ % of predicted <80% and FEV_1_/FVC > 0.7(27). COPD blood gene expression subtypes were previously defined by Chang et al. using network-based stratification (NBS)(24). Of the four NBS subtypes identified in the original publication, the two most prevalent subtypes were analyzed. These subtypes are referred to as the less impaired lung function (LI-NBS) and the more impaired lung function (MI-NBS) subtypes. Current smoking status, inhaled corticosteroid use, and oral corticosteroid use were ascertained by questionnaire. In ECLIPSE, only 8 subjects reported using oral steroids at baseline, and these were removed from subsequent analyses.

### Association of CBC Cell Types with COPD GOLD Stage, Cross-Sectional FEV_1_, and Prospective Change in FEV_1_

Using 4,558 subjects with complete CBC and spirometric data from the COPDGene Phase 2 visit, we plotted the distribution of neutrophil and lymphocyte counts and proportions against GOLD Stage after removing outlying cell count observations greater than +/- 4 SD from the mean. We tested for univariate association between individual cell counts and proportions with post-bronchodilator FEV_1_ % of predicted using Wald tests from linear regression models, and we constructed multivariate regression models relating cell counts and proportions to FEV_1_ adjusting for current smoking status and oral and inhaled steroid use reported at baseline.

In 1,741 smokers from the ECLIPSE Study with complete covariate, CBC, baseline and prospective FEV_1_ measurements, we related absolute cell counts and proportions to post-bronchodilator FEV_1_ % of predicted levels as above. For the analysis of three-year change in FEV_1_ % of predicted levels, we calculated the difference between the first and last available post-bronchodilator FEV_1_ % of predicted measurement in all study subjects (calculated as last measurement - first measurement, i.e. negative values represent decline in lung function). To determine the association between CBC measurements and change in FEV_1_, we used multivariate linear regression models with change in FEV_1_ as the response variable adjusting for baseline FEV_1_, days of follow up, inhaled corticosteroid use at baseline, and smoking status at the first and last study visit. Smoking status was represented in four groups, i.e. current smokers at first and last visit, former smokers at first and last visit, current smoker at first visit and former smoker at last visit, and former smoker at first visit and current smoker at last visit. Models were constructed to analyze cell types individually as well as in the presence of other cell types in the same model. Subjects with less than 1000 days between their first and last spirometric measurements were excluded from analysis.

### Gene Expression

Sample preparation and quality control procedures for genome-wide gene expression data in ECLIPSE have been previously described(28). Standard quality control and quantile normalization were performed. Gene expression data are accessible via GEO [ECLIPSE GSE76705].

### Cell Type Deconvolution and Association of Inferred Cell Types with COPD Status

Cell type deconvolution was performed in 221 ECLIPSE subjects with complete genome-wide gene expression and covariate data. Cell type proportions were inferred using two methods - CIBERSORT(21) and the method of Abbas et al. using linear least squares regression (LR)(20). Cell type reference expression profiles were used from the LM22 pure-cell dataset obtained on 12/21/2015 from the CIBERSORT website (<u>https://cibersort.stanford.edu)</u>. Detailed description is provided in the Online Supplement.

After obtaining cell type estimates of the 22 cell types from both methods, we organized groups of similar cell types into broader categories to create estimates of an additional 9 aggregated groups: CD4+ cells, T cells, B cells, lymphocytes, and monocytes/macrophages. We performed this aggregation by summing individual cell type values for cell types within each category. The 22 inferred cell type proportions and aggregated cell type estimates were tested for association with COPD status and COPD molecular subtypes using the Wilcoxon-Mann-Whitney test. Significant cell type associations were considered to be those with a Wilcoxon-Mann-Whitney test p-val < 0.05 for both CIBERSORT and LR.

### Prediction Models for COPD Status and COPD Molecular Subtypes

Classification of subjects according to COPD status or NBS molecular subtype using estimated cell-type quantities, CBC quantities, and clinical covariates was performed in 221 subjects from ECLIPSE using the support vector machine implementation in the e1071 package(29). Validation within ECLIPSE involved performing one round of partitioning in which half of the subjects were used in the training set and the other half were used in the validation set. Probabilities were returned from the SVM and used with the R package ROCR to generate ROC plots and calculate AUCs(30).

Additional details regarding study cohorts and statistical methods are included in the Supplemental Materials.

## Results

### Relating Circulating Immune Cells to Cross-sectional FEV_1_ and Current Smoking

We examined complete blood count (CBC) data from 4,558 smokers from the COPDGene phase 2 visit and an additional 1,741 smokers with >1000 days of spirometric follow-up data in the ECLIPSE Study. The clinical characteristics and cell type distributions of analyzed subjects in both studies are shown in Table 1. The CONSORT diagram for the analyses of cross-sectional and longitudinal data in ECLIPSE is shown in Supplemental Figure 1, and comparison of characteristics of ECLIPSE analyzed and excluded subjects is shown in Table E1.

**Figure 1:**
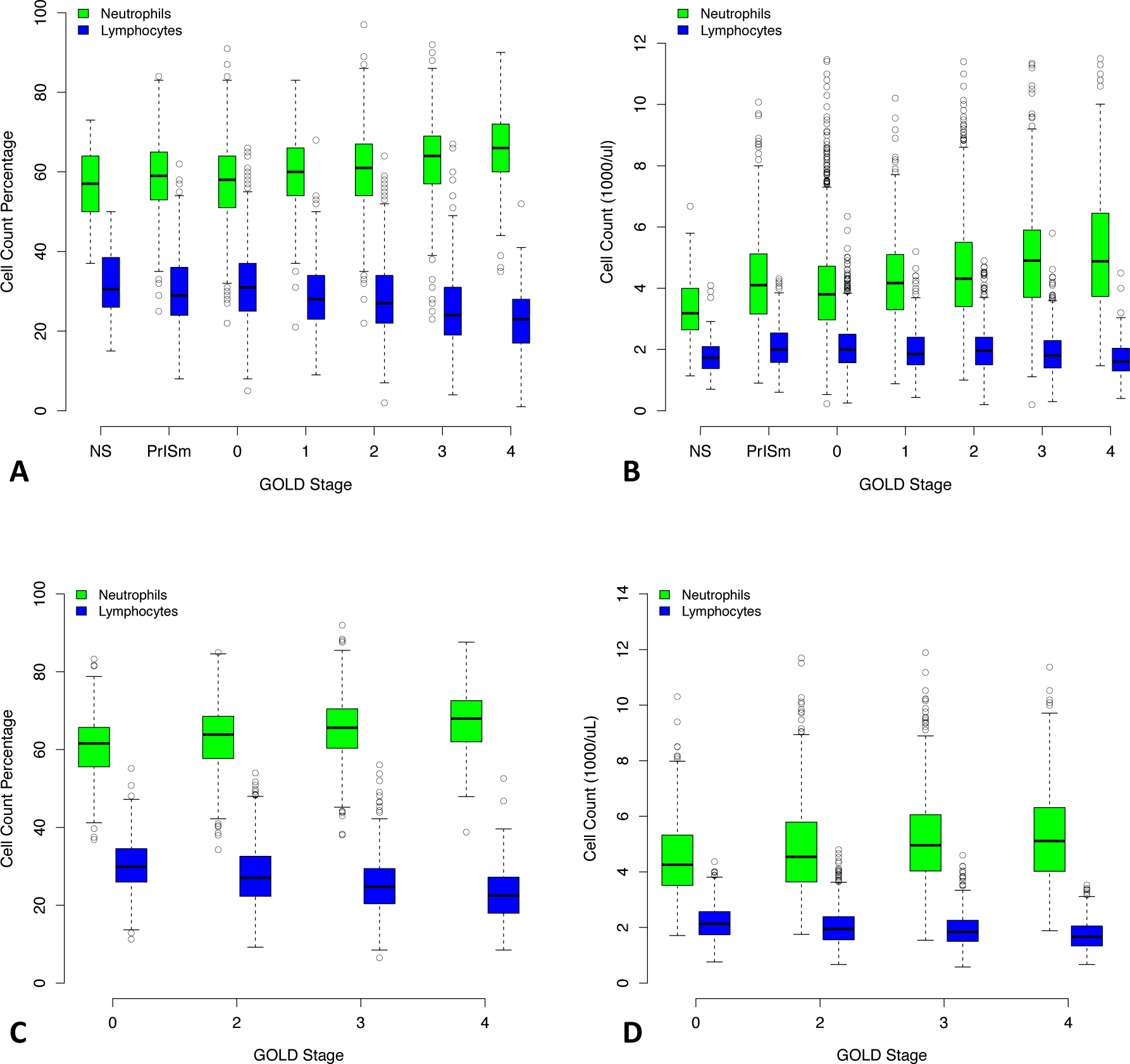
Neutrophil and Lymphocyte Counts and Proportions Stratified by GOLD Spirometric Stage. As GOLD Stage increases, the relative proportions of peripheral neutrophils and lymphocytes increase and decrease, respectively (Panel A, COPDGene and Panel C, ECLIPSE). This phenomenon is driven by an increase in absolute neutrophil count, while the absolute amount of lymphocytes remains relatively stable across GOLD Stages (Panel B, COPDGene and Panel D, ECLIPSE). NS = non-smoker. PRISm = Preserved ratio impaired spirometry, i.e. subjects with FEV_1_/FVC > 0.7 but FEV_1_ % predicted < 80%.

**Table 1.** Characteristics of Analyzed Subjects in COPDGene and ECLIPSE.

|  | COPDGene | ECLIPSE |
| --- | --- | --- |
| N | 4,558 | 1741 |
| Age | 65.5 (8.7) | 61.9 (7.9) |
| Gender, % Female | 50 | 36 |
| Race, % African-American | 27 | 0 |
| FEV1 % of predicted | 78.3 (24.9) | 55.0 (26.1) |
| FEV1/FVC | 0.67 (0.15) | 0.66 (0.21) |
| COPD, % GOLD 2-4 | 35 | 84 |
| Pack Years | 44.0 (24.0) | 45.4 (26.5) |
| Current Smoking, % | 37 | 39 |
| BMI | 28.9 (6.3) | 26.6 (5.3) |
| Oral Steroids, % | 2 | 0 |
| Inhaled Steroids, % | 24 | 59 |
| neutrophil, % | 59.4 (10.0) | 63.9 (8.2) |
| neutrophil, 1000 cells/ul | 4.3 (1.6) | 5.0 (1.6) |
| lymphocyte, % | 29.4 (9.4) | 26.8 (7.6) |
| lymphocyte, 1000 cells/ul | 2.0 (0.7) | 2.0 (0.6) |
| monocyte, % | 8.1 (2.4) | 6.2 (2.1) |
| monocyte, 1000 cells/ul | 0.6 (0.2) | 0.5 (0.2) |
| eosinophil, % | 2.6 (1.7) | 2.8 (1.7) |
| eosinophil, 1000 cells/ul | 0.2 (0.1) | 0.2 (0.1) |
| basophil, % | 0.6 (0.5) | 0.3 (0.2) |
| basophil, 1000 cells/ul | 0.03 (0.04) | 0.03 (0.02) |
| Data are mean (SE) unless otherwise indicated. |  |  |

Linear regression relating the absolute amount and percentage of five cell types to FEV_1_ % of predicted indicated that neutrophils, lymphocytes, monocytes, and eosinophils are strongly correlated with FEV_1_, and there are differences in the pattern of association between absolute counts and cell proportions with COPD severity (Tables 2 and E2). Boxplots showing the amount of each cell type by GOLD stage for COPDGene and ECLIPSE are shown in Supplemental Figures 2 and 3.

**Table 2.**
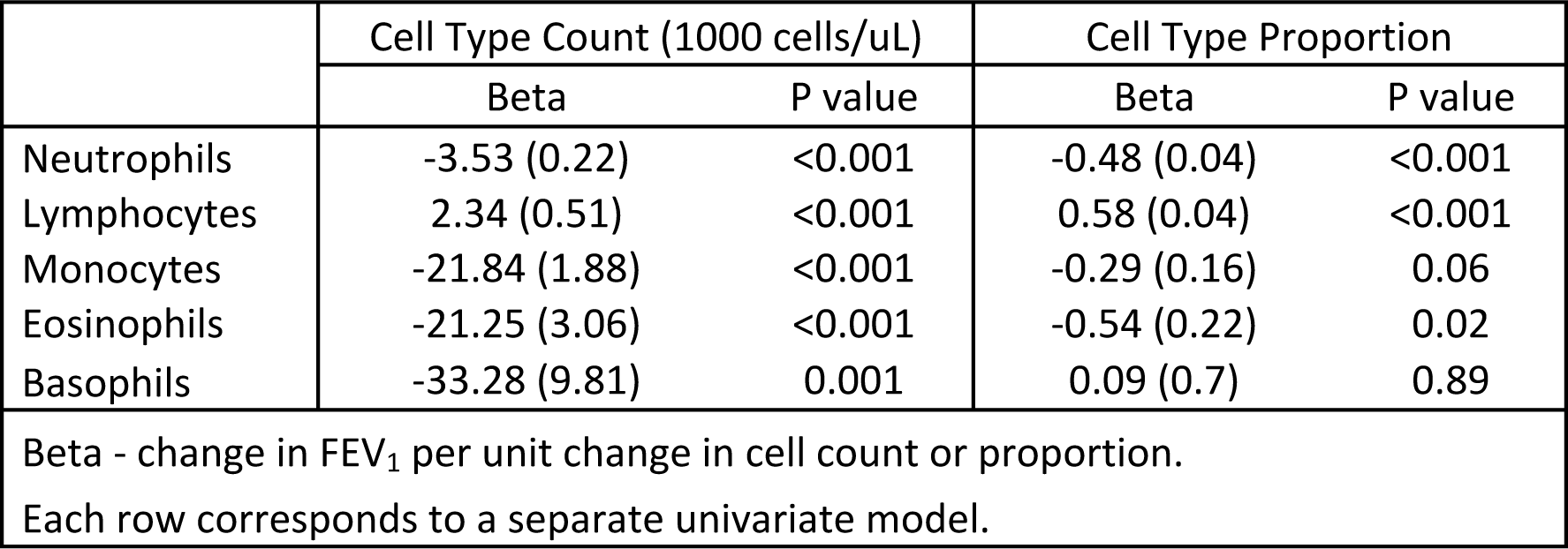
Relationship of Cell Type Counts and Proportions to FEV_1_ % of Predicted in 4,558 Smokers in COPDGene.

|  | Cell Type Count (1000 cells/uL) |  | Cell Type Proportion |  |
| --- | --- | --- | --- | --- |
|  | Beta | P value | Beta | P value |
| Neutrophils | -3.53 (0.22) | <0.001 | -0.48 (0.04) | <0.001 |
| Lymphocytes | 2.34 (0.51) | <0.001 | 0.58 (0.04) | <0.001 |
| Monocytes | -21.84 (1.88) | <0.001 | -0.29 (0.16) | 0.06 |
| Eosinophils | -21.25 (3.06) | <0.001 | -0.54 (0.22) | 0.02 |
| Basophils | -33.28 (9.81) | 0.001 | 0.09 (0.7) | 0.89 |
| Beta - change in FEV<sub>1</sub> per unit change in cell count or proportion. |  |  |  |  |
| Each row corresponds to a separate univariate model. |  |  |  |  |

**Figure 2.**
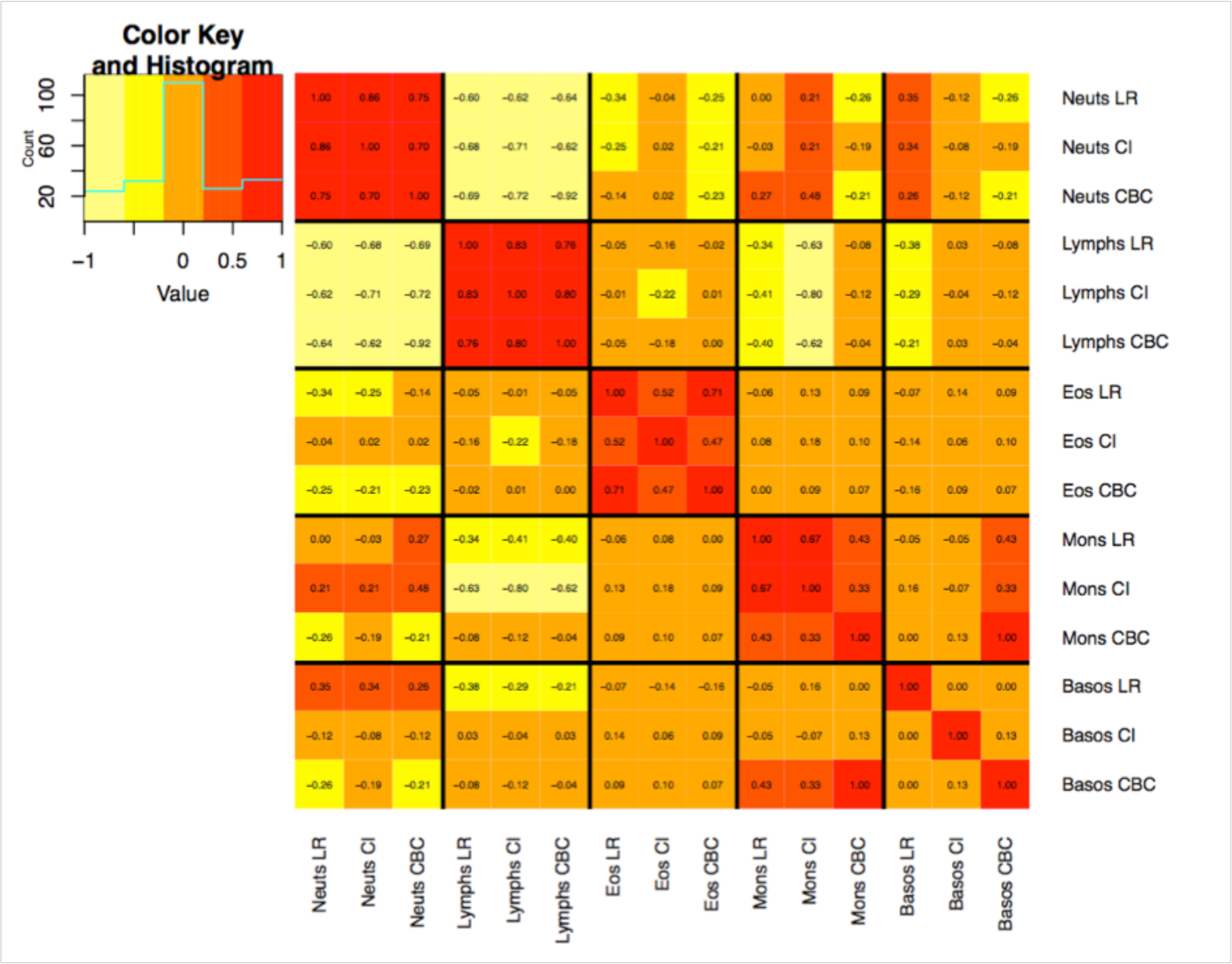
Correlation Between Inferred Cell Subpopulation Proportions and Complete Blood Count Cell Type Proportions. Spearman correlation between estimated cell subpopulations proportions from two deconvolution methods and CBC proportions. Abbreviations: LR-Linear Regression. CI-CIBERSORT. CBC-Complete Blood Count. Neuts-Neutrophils. Lymph-Lymphocytes. Eos-Eosinophils. Mons-Monocytes. Basos-Basophils.

**Figure 3.**
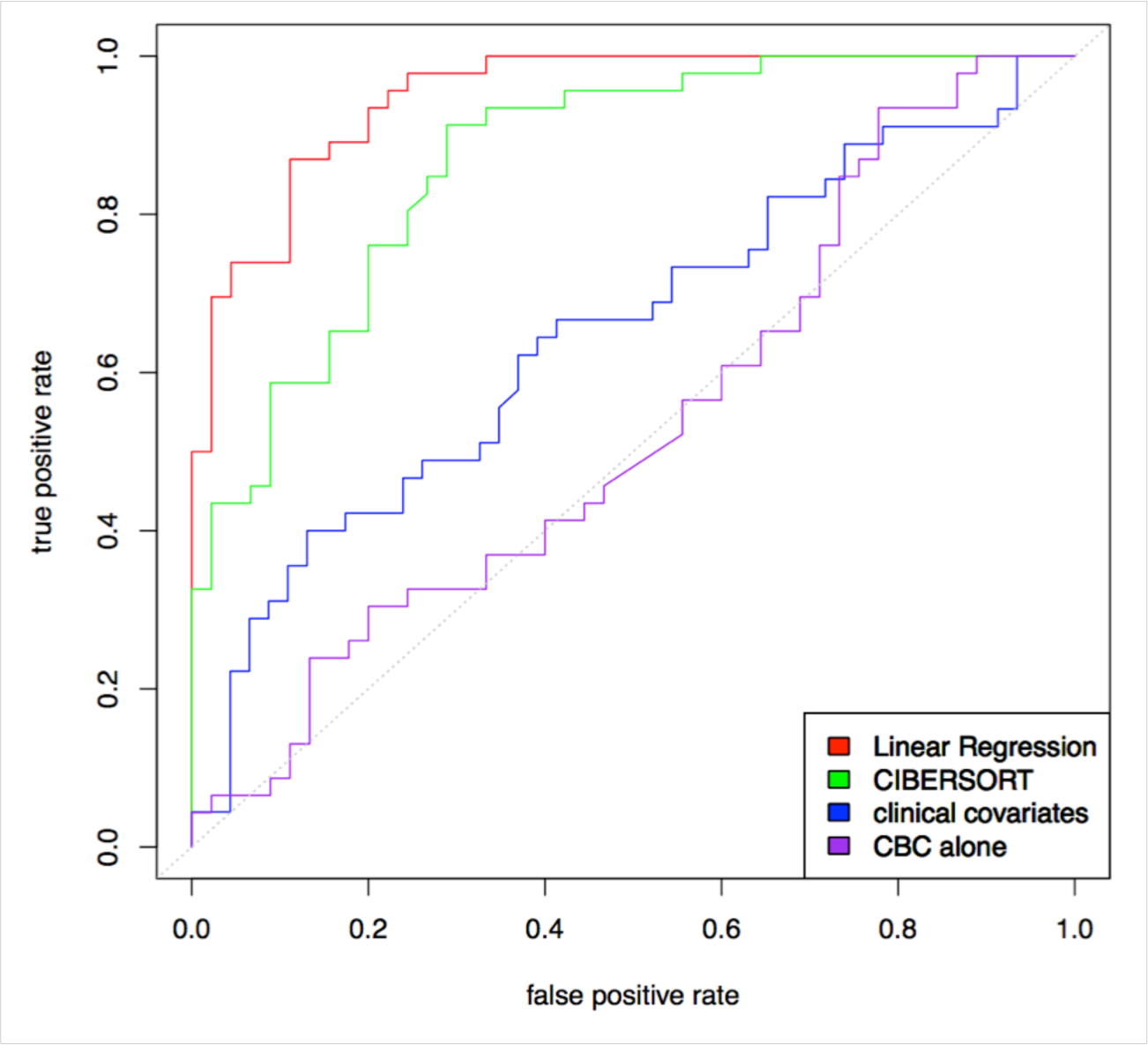
Performance of Predictive Models for COPD Molecular Subtypes Using Complete Blood Counts, Inferred cell Subpopulation Proportions, and Clinical Covariates. Receiver operating characteristic curves demonstrate the predictive performance of SVM classifiers using complete blood count data, inferred cell subpopulation data, and clinical covariates for COPD molecular subtypes in 221 ECLIPSE subjects. Clinical covariates are age, sex, and pack-years.

Given the predominance of neutrophils and lymphocytes in blood, we examined the absolute counts and percentages of these cell types across GOLD stages, and we observed two phenomena. First, with increasing COPD severity, the proportion of neutrophils increases and lymphocytes decreases. However, in terms of absolute cell counts the number of neutrophils increases while the total number of lymphocytes remains relatively stable (Figure 1, Panels A and C), suggesting that the observed changes in neutrophil and lymphocyte proportions associated with COPD severity are primarily driven by an increase in the number of circulating neutrophils. The same pattern is present in ECLIPSE subjects (Figure 1, Panels B and D).

We evaluated these relationships in a series of models in COPDGene relating cell count, cell proportion, and current smoking (CS) to FEV_1_ % of predicted while adjusting for inhaled and oral steroid use (Table 3). The models explaining the largest proportion of variance in FEV_1_, after adjusting for model complexity, included both cell counts and proportions demonstrating that both measures have independent association to FEV_1_. Both lymphocyte and neutrophil absolute counts and percentages were significantly associated with FEV_1_ across most models. The addition of monocyte counts and proportions to the models did not affect the association between neutrophil quantifications and FEV_1_ (data not shown).

**Table 3.** Multivariate Models of the Relationship between FEV_1_ % of Predicted and Neutrophil and Lymphocyte Quantifications.

|  | Neutrophil<br>Count | Neutrophil<br>% | Lymphocyte<br>Count | Lymphocyte<br>% | Current<br>Smoking | Neutrophil<br>Count x<br>Current<br>Smoking | Neutrophil % x<br>Current Smoking | Lymphocyte<br>Count x<br>Current<br>Smoking | Lymphocyte<br>% x Current<br>Smoking | Adjusted<br>R2 |
| --- | --- | --- | --- | --- | --- | --- | --- | --- | --- | --- |
| Cell Proportions | --- | 0.39 (0.10) * | --- | 0.75 (0.11) * | --- | --- | --- | --- | --- | 0.274 |
| Cell Counts | -2.19 (0.20) * | --- | 1.72 (0.44) * | --- | --- | --- | --- | --- | --- | 0.275 |
| Cell Counts +<br>Current Smoking | -2.73 (0.27) * | --- | 1.79 (0.58) * | --- | -5.74 (2.48) * | 1.21 (0.40) * | --- | -0.06 (0.90) | --- | 0.276 |
| Cell Proportions +<br>Counts + Current<br>Smoking | -2.47 (0.62) * | 0.74 (0.14) * | -0.41 (1.32) | 0.94 (0.16) * | -10.99 (18.91) | 0.24 (1.03) | 0.13 (0.23) | 1.85 (2.07) | -0.1 (0.27) | 0.285 |

\* p<0.05
Adjusted R2 - variance explained adjusted for model complexity. Cell counts are in units of 1000 cells/ul.
Each row corresponds to a separate multivariate model. Each model is also adjusted for oral and inhaled steroid use.

### Relating Circulating Immune Cells to Prospective, Three-Year Change in Lung Function

Since the CBC data in COPDGene were obtained at visit 2, longitudinal FEV_1_ analysis measures for this cohort were not available. We performed longitudinal analysis for three-year change in FEV_1_ % of predicted in 1,741 smokers from the ECLIPSE Study who were not taking oral steroids at baseline. In an analysis of single cell type measures, lymphocyte, monocyte, and eosinophil counts and proportions were significantly associated with change in FEV_1_ (Table 4). Higher monocyte levels at baseline were associated with greater FEV_1_ decline, and the opposite pattern was observed for eosinophils. Neutrophil proportions, but not counts, were significantly associated with lung function decline. Larger neutrophil proportions were associated with more lung function decline, with the opposite relationship observed for lymphocyte proportion.

**Table 4.**
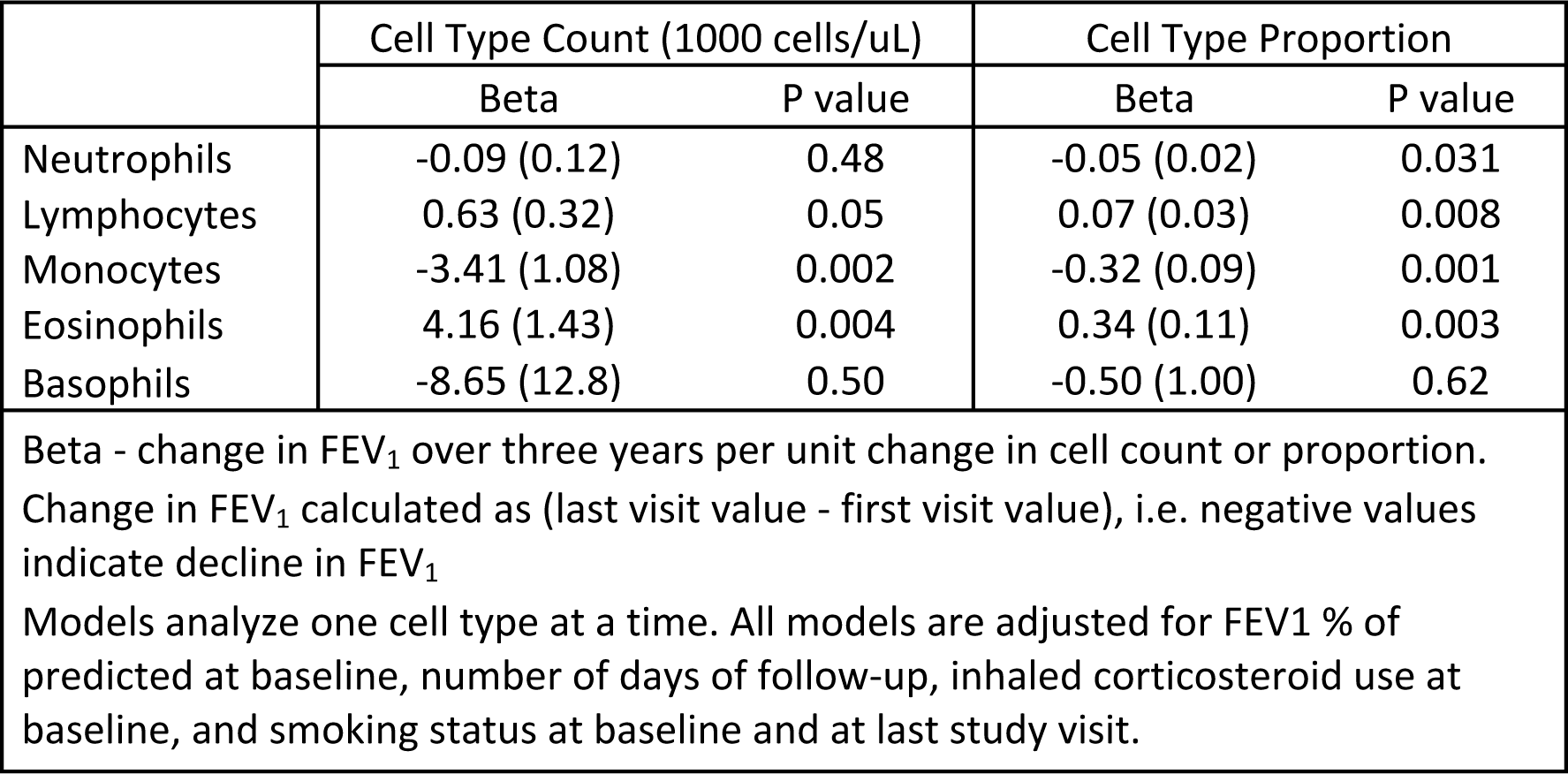
Relationship of Cell Type Counts and Proportions at Baseline to Three-Year Change in FEV_1_ % of Predicted in 1,741 Smokers in ECLIPSE.

|  | Cell Type Count (1000 cells/uL) |  | Cell Type Proportion |  |
| --- | --- | --- | --- | --- |
|  | Beta | P value | Beta | P value |
| Neutrophils | -0.09 (0.12) | 0.48 | -0.05 (0.02) | 0.031 |
| Lymphocytes | 0.63 (0.32) | 0.05 | 0.07 (0.03) | 0.008 |
| Monocytes | -3.41 (1.08) | 0.002 | -0.32 (0.09) | 0.001 |
| Eosinophils | 4.16 (1.43) | 0.004 | 0.34 (0.11) | 0.003 |
| Basophils | -8.65 (12.8) | 0.50 | -0.50 (1.00) | 0.62 |
| Beta - change in FEV<sub>1</sub> over three years per unit change in cell count or proportion.<br>Change in FEV<sub>1</sub> calculated as (last visit value - first visit value), i.e. negative values indicate decline in FEV<sub>1</sub><br>Models analyze one cell type at a time. All models are adjusted for FEV1 % of predicted at baseline, number of days of follow-up, inhaled corticosteroid use at baseline, and smoking status at baseline and at last study visit. |  |  |  |  |

Table 5 demonstrates that for multivariate models including counts and proportions of these four cell types, cell counts, but not proportions, showed significant associations to change in FEV_1_. Absolute counts of monocytes, eosinophils, and lymphocytes were significantly associated with FEV1 decline (p=0.0003, 0.0004, and p=0.02, respectively). Higher levels of monocytes were associated with larger amounts of FEV_1_ decline, and the opposite pattern was present for lymphocytes and eosinophils.

**Table 5.**
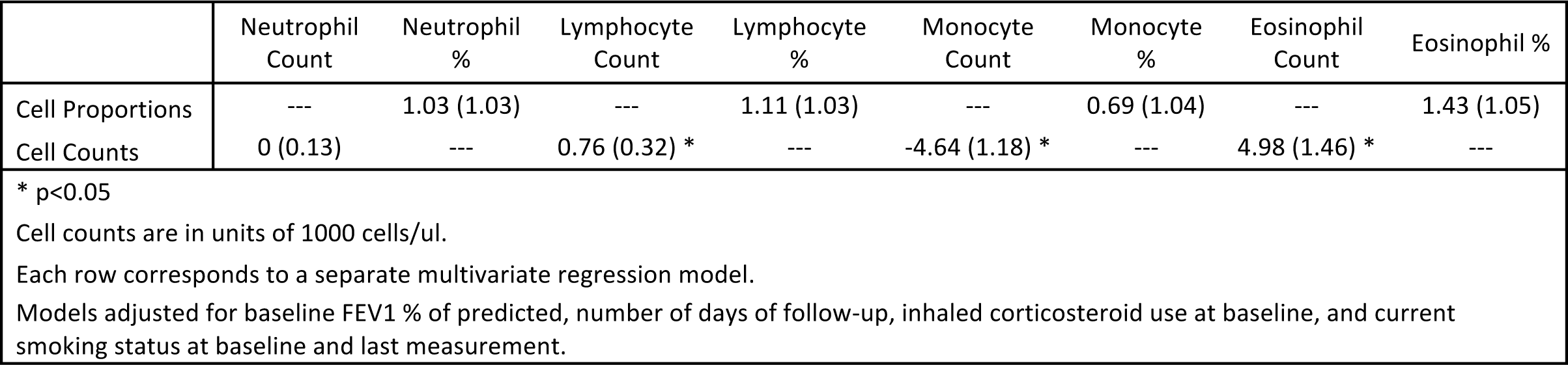
Multivariate Models Relating Cell Type Counts and Proportions at Baseline to Three-Year Change in FEV_1_ % of Predicted in 1,741 Smokers in ECLIPSE.

### Association of Inferred Lymphocyte Subpopulations to COPD and COPD Subtypes

In a subset of 221 subjects from ECLIPSE with complete genome-wide blood gene expression and covariate data (subject characteristics shown in Table E3), we used cell type deconvolution to estimate the proportion of immune cell subpopulations in each study subject, and we related these proportions to COPD case-control status.

To first assess the performance of cell type deconvolution in blood gene expression from smokers, we examined results from applying two methods that have been previously validated for the detection of immune cell types, CIBERSORT and the linear regression (LR) method of Abbas (20, 21). To benchmark these algorithms against known cell type quantifications, we compared their neutrophil, aggregated lymphocyte, aggregated monocyte, eosinophil, and basophil quantifications against concurrently drawn CBC counts (Figure 2). Both methods showed high correlation to neutrophils and lymphocytes (Spearman *r* ranges from 0.7-0.8, p<0.001), with weaker correlations for eosinophils and monocytes. Correlation with basophils was low for both methods. For the inferred proportions of neutrophils and lymphocytes, the correlation between methods was high (0.86 and 0.83, respectively).

We compared the inferred cell type proportions by COPD case/control status and observed that, relative to smoker controls, subjects with COPD had significantly lower levels of aggregated lymphocytes, aggregated T-cells, CD4+ resting memory cells, naive B-cells, and increased levels of monocytes (Table 6).

**Table 6.**
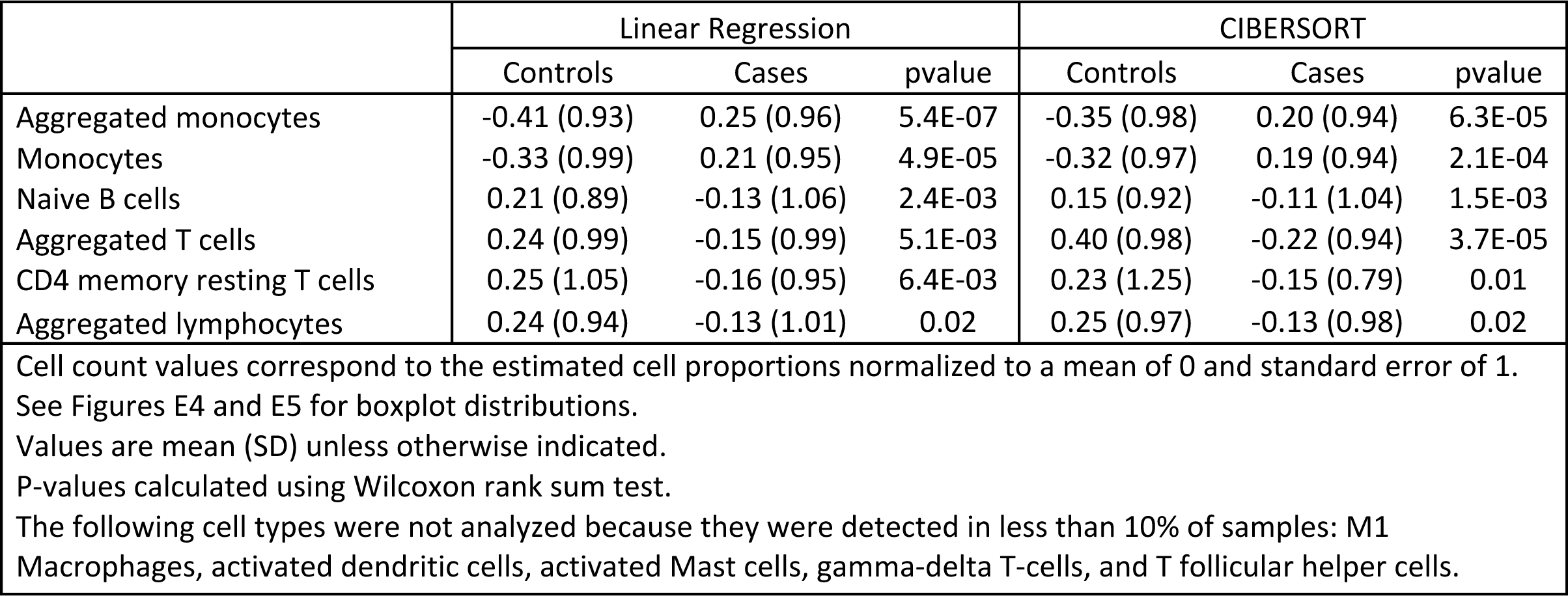
Inferred Immune Cell Types Significantly Associated with COPD Status.

### Inferred Cell Type Proportions Predict COPD Blood Gene Expression Subtypes

In a previous publication, the ECLIPSE blood gene expression data had been used to define COPD molecular subtypes that differed in clinical characteristics and blood gene expression patterns, and we previously demonstrated that these subtypes could not be recovered using CBC data alone(24). To determine whether these molecular subgroups can be accurately predicted from inferred cell count proportions, we trained support vector machine classifiers to predict NBS subtype and COPD case/control status using CBC data, clinical covariates, or inferred cell type proportions. Figure 3 demonstrates that predictive models for NBS subtypes using inferred cell type proportions classified subjects by COPD molecular subtype with high accuracy (AUC = 0.95) and demonstrated better performance than models using only CBC cell-type quantities (AUC = 0.53) or clinical covariates (AUC = 0.65). Predictive models for COPD case/control status using inferred cell type proportions also showed statistically significant, but less powerful, predictive performance (AUC = 0.71), with the cell type subpopulation models still outperforming the models using CBC data (Table E4).

We compared levels of the inferred cell type proportions between the NBS-MI and NBS-LI COPD molecular subtypes and observed that the list of immune cell types that were significantly different between COPD molecular subtypes was more extensive than between COPD cases and controls. This list included T-regulatory cells, CD4 resting and activated memory T-cells, memory B-cells, aggregated T-cells, aggregated B-cells, dendritic cells, and monocytes. (Table E5, Figures E4 and E5).

## Discussion

Using two large cohorts enriched for subjects with COPD, we characterized alterations in circulating immune cell types associated with cross-sectional FEV_1_ and longitudinal FEV_1_ decline. The main findings are: 1) the predominant peripheral immune cell type alteration associated with increasing COPD severity is an increase in the absolute count of neutrophils, 2) monocytes and eosinophils have strong multivariate associations to prospective change in FEV_1_, and 3) cell-type estimates from gene expression deconvolution methods show good accuracy for some cell types.

Prior immunologic studies of COPD have identified important associations with increased innate immune activation and progression of COPD including neutrophil stimulation(31, 32), alveolar macrophage immune surveillance(33), protease/matrikine activation(34), and activation of the dendritic cell/macrophage axis(35). Most of these studies have been performed in murine models or small to moderate sized study samples with limited ability to control for the effects of current smoking. Our findings complement and extend previous results by demonstrating that 1) the decrease in overall lymphocyte proportion in COPD is primarily driven by an increase in absolute neutrophil counts, 2) absolute counts and proportions of immune cell types have independent, statistically significant associations to lung function, and 3) absolute monocyte and eosinophil counts are predictive of COPD disease progression. This point extends previous observations relating total peripheral leucocyte count to cross-sectional and longitudinal lung function(16-18) by implicating specific myeloid cell types. The fact that monocytes were associated with cross-sectional FEV_1_ and prospective FEV_1_ decline, whereas neutrophils only showed multivariate association to cross-section FEV_1_ is an interesting finding. This suggests that increased circulating monocytes may play an important role in initiating or maintaining the inflammatory processes responsible for ongoing lung destruction. While circulating neutrophils are clearly associated with COPD severity, they were not an independent predictor of decline after accounting for other cell types and covariates. However, precise mechanistic hypotheses are beyond the scope of this epidemiologic study and would require detailed assessment of both the lung and systemic compartments.

To study immune cell subpopulations not quantified by CBC, we explored the use of cell type deconvolution to quantify 22 distinct cell subpopulations in a subset of ECLIPSE subjects with blood genome-wide gene expression data. These methods are a promising alternative for estimating cell type proportions in large study samples with available expression data. However, these approaches have not been widely applied in smokers with COPD. Our findings demonstrate that in smokers enriched for COPD, the deconvolution approaches studied yielded consistent and reasonably reliable estimates of neutrophil and lymphocyte cell proportions with mixed performance for other cell types. Inferred cell type proportions enabled significantly better prediction of externally defined COPD molecular subtypes than CBC and clinical data alone, providing indirect evidence that these inferred proportions capture meaningful information on the cell type composition of bulk blood expression samples. These data provide proof-of-concept of the feasibility of using cell type deconvolution to study immune cell subpopulations in large cohorts of smokers with available blood gene expression data.

Analysis of the cell type deconvolution data supports previous observations of an overall decrease in lymphocytes and T-cells in the COPD state, coupled with an increase in monocytes. When we studied deconvolved cell types in previously defined COPD molecular subtypes, differences in cell type proportions were more pronounced, with the more severely affected subtype characterized by increased monocytes, T-regulatory cells, memory T-cells, and memory B-cells as well as decreased total lymphocytes. This pattern is consistent with the expected behavior of T-regulatory cells, which play a critical immunomodulatory role by suppressing other lymphocyte populations in part through IL-10 and TGF-β signalling(36). Overall, these findings provide additional support for the model that the circulating immune response to CS and COPD is characterized by distinct aspects of suppression of the adaptive immune response and a chronic increase in myeloid cell types, and it suggests that within smokers there are patterns of coordinated immune response that can be used to identify clinically distinct subgroups of subjects.

The main strength of this study is that peripheral cell type quantifications were available from a large number of smokers with a broad range of lung function from two independent studies. The study design also enabled the study of immune cell alterations adjusting for CS, a major confounder of the relationship between immune cell alterations and COPD severity. The use of cell type deconvolution to study the immune response in COPD is novel and enabled the simultaneous study of a large number of lymphocyte subpopulations. Because CBC quantifications and blood gene expression were available in the same ECLIPSE subjects from the same time point, we could benchmark our deconvolution approaches against a known standard.

This study also has important limitations. We did not have access to immune cells in the lung or specific lung compartments in our study subjects, thus we could not relate the blood observations to the lung compartment. We also were not able to characterize immune cell functional states through cytokine profiling or quantification of response to antigenic stimulation. While we observed significant associations between circulating immune cell subpopulations and COPD, further study is required to determine the pathophysiological significance of these observations. We did not have flow cytometry values against which we could benchmark our cell type deconvolution estimates, but we did have Coulter counter data available, and our deconvolution results were generated using methods that has been previously validated against flow cytometry for immune cell populations(20, 21).

In conclusion, analysis of CBC counts and proportions in >6,000 subjects from the COPDGene and ECLIPSE studies demonstrated that cross-sectional FEV_1_ is associated with alterations in multiple circulating immune cell types, including total neutrophil count. COPD disease progression, as quantified by decline in FEV_1_, is associated with increased absolute monocyte counts and decreased lymphocyte and eosinophil counts at baseline. Cell type deconvolution is a viable approach to simultaneously study multiple immune cell populations in smokers with COPD. Future studies to characterize COPD-related alterations in more fine-grained immune cell types will benefit from quantification of both cell type proportions and absolute counts.

## Authors’ contributions

### Study concept and design

*Stromberg, Castaldi, Bowler*

### Acquisition, analysis, or interpretation of data

*Stromberg, Castaldi, Bowler*

### Drafting of the manuscript

*Stromberg, Castaldi, Bowler*

### Critical revision of the manuscript for important intellectual content

*All authors*

### Statistical analysis

*Stromberg, Castaldi, Bowler*

### Obtained funding

*Castaldi and Bowler*

## Declarations

### Ethics approval and consent to participate

For both COPDGene and ECLIPSE, the institutional review boards of all participating centers approved these studies, and written informed consent was obtained from all subjects.

### Consent for publication

Not applicable.

### Availability of data and material

Gene expression data used in this study are accessible via GEO [ECLIPSE GSE76705, COPDGene GSE42057].

